# A novel *Parvimonas* OTU is inversely associated with having caries or fillings in children

**DOI:** 10.64898/2025.12.16.694580

**Authors:** Veera F. Kukkonen, Nitin Agrawal, Anna Liisa Suominen, Aino-Maija Eloranta, Matti Närhi, Jukka Leinonen, Pirkko Pussinen, Timo A. Lakka, Heli Viljakainen

**Affiliations:** Institute of Dentistry, Faculty of Health Sciences, University of Eastern Finland, Kuopio, Finland; Folkhälsan Research Center, Helsinki, Finland; Faculty of Medicine, University of Helsinki, Helsinki, Finland; Oral and Maxillofacial Teaching Unit, Primary Oral Health Care Services, Wellbeing Services County of North Savo, Kuopio, Finland; Institute of Biomedicine, School of Medicine, University of Eastern Finland, Kuopio, Finland; Institute of Public Health and Clinical Nutrition, School of Medicine, University of Eastern Finland, Kuopio, Finland; Department of Medicine, Endocrinology and Clinical Nutrition, Kuopio University Hospital, Wellbeing Services County of North Savo, Kuopio, Finland; Department of Oral and Maxillofacial Diseases, University of Helsinki, Helsinki, Finland; Department of Clinical Physiology and Nuclear Medicine, Kuopio University Hospital, Kuopio, Finland; Foundation for Research in Health Exercise and Nutrition, Kuopio Research Institute of Exercise Medicine, Kuopio, Finland

**Author notes:** Equal contribution (Veera F. Kukkonen, Nitin Agrawal). Equal contribution (Timo A. Lakka, Heli Viljakainen). **Corresponding author**: Timo Lakka, Professor of Medical Physiology, Internist, Institute of Biomedicine, School of Medicine, University of Eastern Finland, Kuopio Campus, Yliopistonrinne 3, 70211 Kuopio, Finland.

**Keywords:** microbiota, saliva, oral health, pedodontics

## Abstract

**Background:** Dental caries is the most common chronic oral disease in children, but its etiology is complex and multifactorial. The association of the oral microbiota with caries is underexplored in children. We investigated saliva microbiota in Finnish children and aimed to identify microbial biomarkers associated with the risk of having caries or fillings (CF).

**Methods:** We examined 400 Finnish children (180 girls, 220 boys) aged 7-9 years who provided stimulated saliva samples and underwent clinical oral examinations at baseline of the Physical Activity and Nutrition in Children study. Of these children, 261 (65%) were with CF and 139 (35%) were without CF. We characterized the saliva microbiota using the 16S rRNA gene sequencing, examined differences in the abundance and composition at the phylum and genus level between these two groups, and studied the associations of selected Operational Taxonomic Units (OTUs) with the risk of having CF in crude and adjusted (OTUs, age, Silness-Löe plaque index) models.

**Results:** Alpha diversity and richness did not differ between the groups (*p*>0.1 for Shannon, inverse Simpson, and Chao1 indices) in the total sample nor when analyzed among girls and boys separately. Beta diversity differed between the groups in the total sample, and when girls and boys were analyzed separately (*p* corrected by false discovery rate method [*p_corr_*] ∼ 0.001). In total, 80 OTUs differed between the groups at the genus level (*p_corr_*<0.01). Two OTUs were independently associated with the risk of having CF. A novel *Parvimonas* OTU was inversely associated with having CF (odds ratio [OR_crude_] 0.49, 95% confidence interval [CI] 0.32 to 0.75). In addition, an OTU of probably *Streptococcus mutans* was positively associated with having CF (OR_crude_ 6.01, 95% CI 3.49 to 10.36).

**Conclusion:** A *Parvimonas* species represented by a novel OTU is independently and inversely associated with the risk of having CF in children.

## Background

Saliva is an exceptional biofluid for studying the oral microbiota, as it contains a variety of bacteria from different oral niches, including the supra- and subgingival plaque, crevicular fluid, and tongue [1]. Saliva microbiota is an emerging field of study especially for population level studies due to its noninvasive nature and ease of collection [2]. Dysbiosis in the oral microbiome has been associated with various oral diseases such as caries in children [1, 3] and adults [4] as well as periodontitis and systemic diseases such as type 2 diabetes in adults [5, 6].

Dental caries is a common childhood oral disease that results from oral bacteria metabolizing easily fermentable carbohydrates to produce acids that demineralize dental hard tissues over time [7–10]. The etiology of caries is biofilm-mediated and multifactorial, and includes physical, biological, environmental, behavioral, and lifestyle-related factors [10] such as saliva secretion, fluoride concentration of drinking water, frequency of consuming easily fermentable carbohydrates [11], level of oral hygiene [12] plus biofilm formation and maturation [10]. Dental biofilm becomes cariogenic due to a maturation process where the biofilm not only increases in thickness and mechanical stability but also the proportion of acidogenic and aciduric microbes increases. These taxa comprise the highly acidogenic *Streptococcus mutans* which has been extensively studied because of its role in caries [13], but also e.g. *Leptotrichia*, *Veillonella*, *Bifidobacterium*, and *Porphyromonas* [14]. In addition, genetic susceptibility may increase the risk of caries [15, 16].

Several studies have shown differences in the saliva microbiota between children and adolescents with and without caries [8, 14, 17–19]. The results, however, are inconsistent, possibly due to limited sample sizes, high inter-individual variation in saliva microbiota, differences in age or sequencing methods, and the role of various oral bacteria in the development of caries. The findings of a systematic review on the associations of oral microbiota with caries in adolescents indicated that multiple bacterial taxa are linked to caries, supporting the concept that caries is a polymicrobial disease rather than the result of a few specific cariogenic species [14]. However, understanding the contribution of specific oral bacteria to the microbiota imbalance is essential in preventing and controlling caries [12, 18, 20, 21]. Many bacteria that were previously undetectable can now be identified cost-effectively using 16S rRNA gene sequencing [22].

Here, we study the diversity and composition of saliva microbiota using 16S rRNA gene sequencing in a relatively large general population of Finnish children aged 7-9 years with a clinically examined caries status to identify microbial OTUs associated with having caries or fillings (CF).

## Methods

### Study design and study population

The present analyses are based on the baseline data from the Physical Activity and Nutrition in Children (PANIC) study, a non-randomized controlled trial in a general population of children from the city of Kuopio, Finland (https://ClinicalTrials.gov <u>NCT01803776</u>) [23]. The PANIC study has been carried out in accordance with the principles of the Declaration of Helsinki as revised in 2008. The study protocol was approved by the Research Ethics Committee of the Hospital District of Northern Savo in 2006 (Statement 69/2006). Written informed consent was received from the caregivers of the children and assent to participation from the children.

Between 2007 and 2009, 736 children aged 7–9 years, starting their first grade in primary schools of Kuopio, were invited to participate in the study. Altogether, 512 (70%) children accepted the invitation and attended the baseline examinations. There was no notable difference in sex distribution, age, height, or body mass index - standard deviation score (BMI – SDS) between the participants and all children who started the first grade in the city of Kuopio in 2007–2009. Six children were excluded due to physical disabilities that could hamper their participation. In addition, two families withdrew their consent. The final study sample comprised 504 children at baseline. Altogether, 487 children underwent a clinical oral health examination. Of these children, 425 had complete oral health data and saliva samples for the present study. After excluding 18 participants with missing information on microbiota and 7 children whose dental information could not be verified, the final study sample included 400 children.

### Oral health examinations

Experienced dentists of the research group performed oral health examinations according to a strict predefined protocol used in oral health examinations offered to all Finnish children. The oral health examination has been explained in detail elsewhere [24]. For the statistical analyses, the children were divided into those with having caries (C) or fillings (F), combined as CF, and those without CF. Prior to baseline examination, no teeth were missing or had been extracted because of caries. The Silness–Löe Plaque index was used to register plaque [25].

### Assessment of body size, oral self-care habits, household income and dietary factors

Body height and weight were assessed by a trained research staff during a study visit after overnight fasting, as detailed elsewhere [24]. BMI was calculated by dividing body weight (kg) with body height (m) squared, and BMI-SDS was calculated using Finnish reference data [26]. The parents and caregivers reported the Household income and tooth brushing frequency in a questionnaire. Tooth brushing frequency was categorized as at least twice a day, once a day, several times a week, once a week or less often, and never, and it was further dichotomized as at least twice a day and less often for statistical analyses. Household income was categorized as ≤30 000 €/y, 30 001–60 000 €/y, and ≥60 001 €/y for the statistical analyses.

Food consumption, eating frequency, and nutrient intake were recorded over a four-day period at baseline, which included either two weekdays and two weekend days, or three weekdays and one weekend day. Clinical nutritionists reviewed the records with families to ensure accuracy [27]. The intake of energy and nutrients, including sugar, vitamins, and other nutrients, from all dietary sources were analysed using the Micro Nutrica^®^ software, Version 2.5 [27, 28].

The overall diet quality was assessed using the Baltic Sea Diet Score (BSDS), which is derived from the habitual dietary intake of the Nordic population [29]. BSDS was calculated by summing scores from six food consumption components and two nutrient intake components, each of which was rated by quartiles among children, as described earlier [28]. Food components included fruits and berries, vegetables (excluding potatoes), low-fat milk, fish, high-fibre grain products, and red meat/sausages. Nutrient components were total fat intake and the polyunsaturated fatty acid to saturated fatty acid ratio [29]. Scores for BSDS ranged between 0 and 24, higher scores indicating better diet quality.

### Saliva sample collection

A trained research staff collected stimulated saliva samples after a 12-hour overnight fasting. The children were asked to wash their hands and to chew a paraffin capsule. During the first 30 seconds, the children spat the saliva into a sample cup, and this specimen was thrown away. The saliva secreted during the next three minutes was collected in a separate sample cup. The children were instructed not to swallow saliva during the saliva collection. The collected saliva was stored frozen at -80 °C until biochemical analyses.

### Amplification and sequencing

A detailed 16S rRNA gene sequencing protocol has been described earlier [30]. In short, bacterial cell lysis and mechanical disruption by bead-beating was performed. The DNA extraction was conducted using the CMG-1035 saliva kit and Chemagic MSM1 nucleic acid extraction robot (PerkinElmer) at the Technology Centre, Sequencing Unit of the Institute for Molecular Medicine Finland. Amplification and sequencing libraries were prepared according to our in-house 16S PCR amplification protocol detailed elsewhere [31]. All samples were analysed together to avoid any technical variation.

### Bioinformatics and biostatistical analysis

Quality filtering and sequence processing were carried out using the CLC Genomics Workbench, Version 22 (https://digitalinsights.qiagen.com) and were aligned to the SILVA 16S rRNA database, Version 138.1, clustered into operational taxonomic units (OTUs) described briefly earlier [31]. In detail, the sequence reads containing ambiguous bases, and more than one mismatch in the primer sequence, less than 40 base-pair assembly overlap or sequences, and over five unaligned mismatched ends under the default parameters in CLC Genomics Workbench were removed to ensure the data quality. Assembled reads with less than 100 base-pairs and more than 470_base-pairs were removed from the analysis. No samples were omitted during the analysis due to low sequencing depth (<10,000 reads). The sequencing generated approximately 50 million reads from the 425 samples. The mean read count per sample was 115,423 (ranging between 12,491 and 155,564). The OTUs which were present less than 20 times (in aggregate; less than 5%) among the 425 samples were filtered out as ‘low counts’. Thus, a total of 2507 OTUs were obtained. Altogether, 25 saliva samples were removed during analysis because of missing or incomplete taxonomic information or metadata, decreasing the total number of saliva samples to 400 (220 boys, 180 girls) and OTUs to 2479. These OTUs were divided into 8 phyla, 12 classes, 29 orders, 45 families, and 87 genera.

Alpha diversity was calculated using the Shannon index, inverse Simpson index, and Chao1 index. Beta diversity, denoting the variation in community composition between the children with or without CF was estimated with the Principal Coordinate Analysis using the Bray-Curtis dissimilarity index and with the Permutational Multivariate Analysis of Variance (PERMANOVA) using the *adonis* function from the *vegan* package with 999 permutations. Due to multiple testing, *p*-values were corrected by the Benjamini-Hochberg false discovery rate method and presented as *p_corr_*.

Relative bacterial composition was calculated at the phylum and genus levels. We identified differentially abundant OTUs at the genus level between groups using general linear models with a negative binomial distribution implemented in the DESeq2 package, Version 1.34.0. Bioinformatic analyses were conducted by the R software, Version 4.0.4, using the *Bioconductor* (Version 3.12.0), *Microbiome* (Version 1.12.0), *Vegan* (Version 2.5-7), and *Phyloseq* (Version 1.34.0) packages. A nucleotide BLAST search (https://blast.ncbi.nlm.nih.gov/Blast.cgi) was performed for the selected OTUs to check for available species-level information.

Backward stepwise logistic regression analysis with Wald method was conducted to choose the determinants for CF. Each OTU was dichotomized (absence and presence of that OTU) and additionally categorized into absence, low abundance (< median for abundance), and high abundance (≥ median for abundance). Logistic regression analysis was applied to analyze the association of each OTU with having CF in both crude and adjusted models. The results were reported as odds ratios (ORs) and their 95% confidence intervals (CIs). These analyses were performed with the IBM SPSS Statistics software, Version 26.0 (IBM Corp., Armonk, NY, USA). In all these analyses, a 5% statistical significance level was adopted.

## Results

### Basic characteristics of children

The mean (SD) age of children was 7.6 (0.4) years, 45% were girls, and 98% were Caucasians. The basic characteristics of the children were similar between the two groups (**Table 1**).

**Table 1:** Basic characteristics of children.

|  | All children<br>(n=400) | with CF (n=261) | without CF (n=139) |
| --- | --- | --- | --- |
| Sex, n (%) |  |  |  |
| Girls | 180 (45.0) | 119 (45.6) | 61 (43.9) |
| Boys | 220 (55.0) | 142 (54.4) | 78 (56.1) |
| Age, years | 7.6 (0.4) | 7.6 (0.4) | 7.6 (0.3) |
| BMI-SDS | -0.16 (1.1) | -0.12 (0.1) | -0.24 (0.1) |
| Silness–Löe Plaque index<sup>1</sup> | 4.2 (0.2) | 4.8 (0.2) | 3.3 (0.2) |
| Tooth brushing frequency, n (%)<sup>2</sup> |  |  |  |
| At least twice a day | 225 (59.4) | 137 (55.9) | 88 (65.7) |
| Less often than twice a day | 154 (40.6) | 108 (44.1) | 46 (34.3) |
| Household income, n (%)<sup>3</sup> |  |  |  |
| ≤ 30 000€/y | 81 (20.6) | 61 (23.8) | 20 (14.5) |
| 30 001€ - 60 000€/y | 168 (42.6) | 107 (41.8) | 61 (44.2) |
| ≥ 60 001€/y | 145 (36.8) | 88 (34.4) | 57 (41.3) |
| Baltic Sea Diet Score<br>(range 0-24)<sup>4</sup> | 11.9 (0.2) | 11.7 (0.3) | 12.3 (0.4) |
| Sucrose intake, E%<sup>5</sup> | 12.8 (3.7) | 12.9 (0.3) | 12.6 (0.3) |

Data are presented as means (standard deviations) for continuous variables and numbers (percentages) for categorical variables.
<sup>1</sup>Data were available for 389 children (173 girls and 216 boys).
<sup>2</sup>Data were available for 379 children (171 girls and 208 boys).
<sup>3</sup>Data were available for 394 children (178 girls and 216 boys).
<sup>4</sup>Data were available for 338 children (156 girls and 182 boys).
<sup>5</sup>Data were available for 338 children (156 girls and 182 boys), and sucrose intake was presented as a percentage of total energy intake.

### Alpha diversity and richness

Alpha diversity indices (Shannon, inverse Simpson, Chao1) did not differ between the sexes, suggesting that primary analyses could be performed by combining data on girls and boys. Alpha diversity (*p*=0.2 for Shannon index, *p*=0.91 for inverse Simpson index), and richness (*p*=0.79 for Chao1 index) did not differ between 261 children with CF and 139 children without CF, either.

### Beta diversity

Beta diversity, estimated with the Principal Coordinate Analysis using the Bray-Curtis dissimilarity index, differed between children with CF and children without CF (*p_corr_*=0.001) (**Figure 1A**). This dissimilarity between the groups persisted when we analyzed girls (*p_corr_*=0.002) and boys (*p_corr_*=0.001) separately.

**Figure 1:**
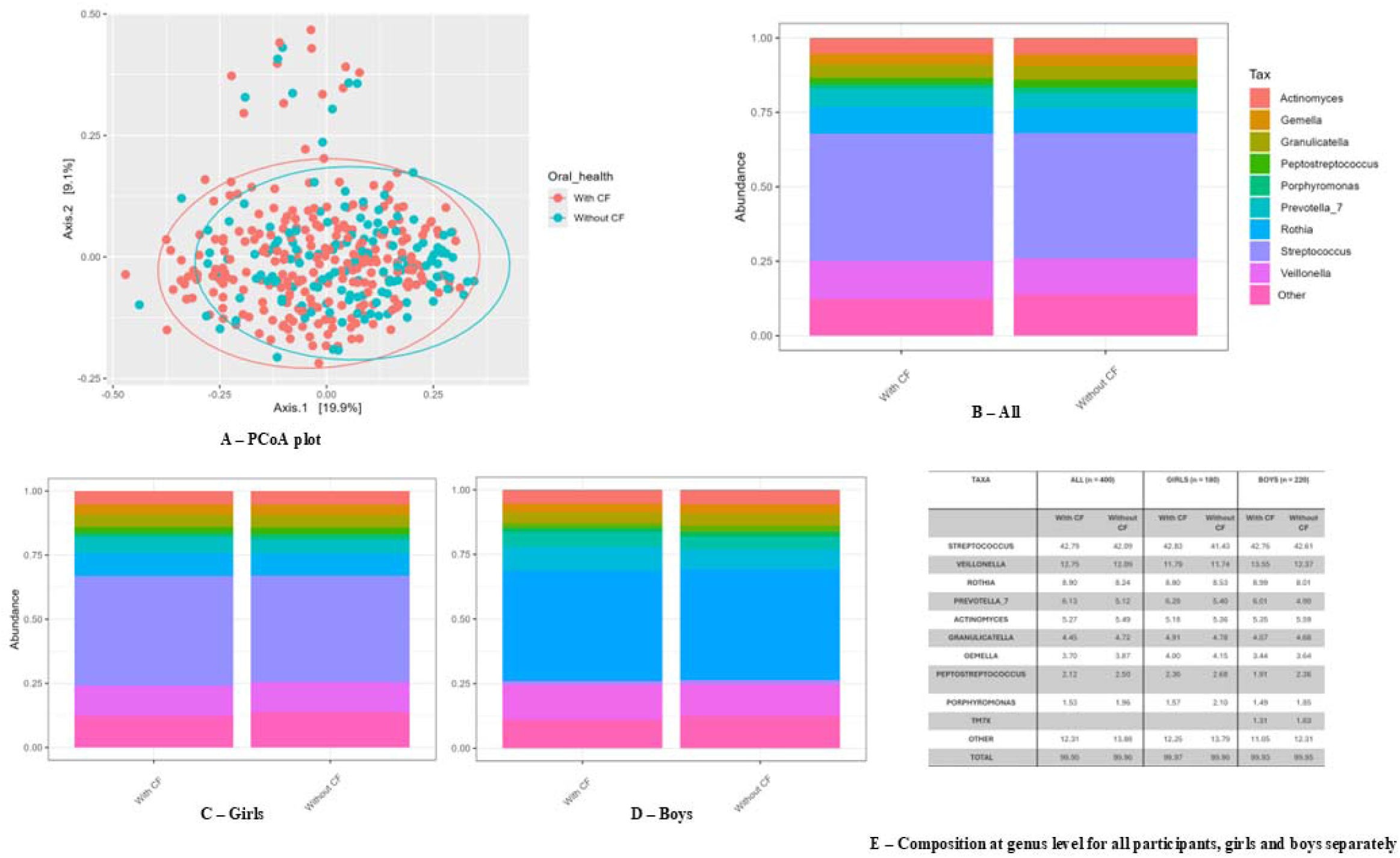
Diversity and composition of the microbiota between children with CF and children without CF. (A) Beta Diversity – Principal Coordinate Analysis (PCoA) plot with Bray-Curtis dissimilarity index between children with CF and children without CF. Abundance at the genus level in (B) all children, (C) girls, and (D) boys. (E) Abundance percentages at the genus level.

### Microbiota abundance

The most common saliva microbiota at the phylum level in children with CF were *Bacillota* (68.8%), *Actinomycetota* (15.8%), *Bacteroidota* (8.9%), and *Pseudomonadota* (2.6%). These proportions were similar in children without CF, and when girls and boys were analyzed separately (**Supplementary Table 1, Supplementary Figure 1**).

At the genus level, the most common saliva microbiota in children with CF were *Streptococcus* (42.7%), *Veillonella* (12.7%), *Rothia* (8.9%), *Prevotella 7* (6.1%), and *Actinomyces* (5.2%) (**Figure 1B, 1E**). These proportions were similar in children without CF. These proportions were also similar between the groups when analyzed for girls and boys separately (**Figure 1C, 1D, 1E**). Interestingly, *TM7x* genera (phylum: Saccharibacteria) was among the 10 most common genera in boys (1.31% in boys with CF, 1.63% in boys without CF) but not in girls nor the whole sample of children.

### Differentially abundant OTUs and their association with caries or fillings

Altogether, 80 OTUs differed between the groups (*p_corr_*<0.01) (**Figure 2**). After filtering out OTUs with low base mean (< 20), one OTU from *Streptococcus*, *Parvimonas*, and *Leptotrichia* genera each that had the largest positive or negative log_2_-fold changes were selected for further analyses **(Table 2)**. The OTU AJ243965.1.1512 within the *Streptococcus* genus had the highest positive log_2_-fold change, indicating an enrichment in children with CF, whereas the OTU FJ470585.1.1487 of *Parvimonas* had the highest negative log_2_-fold changes, indicating a depletion in children with CF. Additionally, the OTU JQ451131.1.1383 of *Leptotrichia* with a negative log_2_-fold change was included in the stepwise analyses.

**Figure 2:**
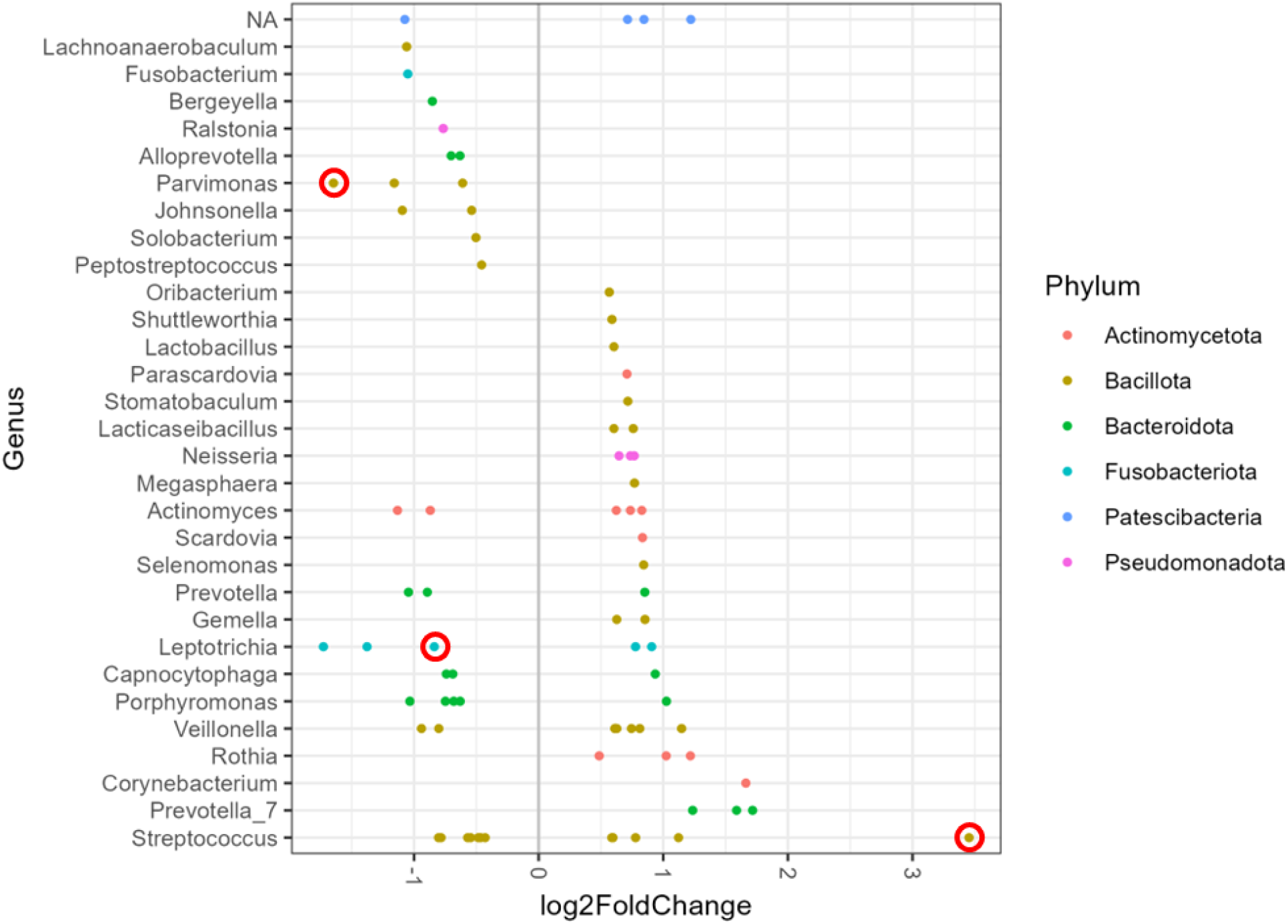
Differentially abundant OTUs (*p_corr_<0.01)* between children with CF and children without CF. Sample-based Log_2_-fold changes (X-axis) on the genus (Y-axis) are shown. The chosen OTUs are circled in red. The positive log_2_-fold change indicates increased risk for CF and the negative log_2_-fold change decreased risk for CF.

**Table 2:** Differentially abundant OTUs in children with or without CF.

| OTU Names | Log<sub>2</sub><br>fold<br>change | *p*-value | $p_{corr}$ | Base<br>Mean | Phylum | Genus |
| --- | --- | --- | --- | --- | --- | --- |
| **AJ243965.1.1512** | **3.45** | **5.95E-39** | **7.31E-36** | **40.71** | **Bacillota** | **Streptococcus** |
| AY281085.1.1430 | -0.42 | 0.0005 | 0.008 | 1282.93 | Bacillota | Streptococcus |
| JX861483.1.1457 | -0.46 | 0.0005 | 0.008 | 832.16 | Bacillota | Streptococcus |
| GU045364.1.1452 | -0.78 | 5.16E-05 | 0.001 | 20.31 | Bacillota | Streptococcus |
| **JQ451131.1.1383** | **-0.83** | **8.72E-05** | **0.002** | **41.91** | **Fusobacteriota** | **Leptotrichia** |
| **FJ470585.1.1487** | **-1.64** | **7.58E-09** | **7.16E-07** | **88.48** | **Bacillota** | **Parvimonas** |

Filtered OTUs with corrected *p*-values <0.01 and base mean > 20, sorted in the descending order of log<sub>2</sub>-fold changes. The OTUs with the highest positive or negative log<sub>2</sub>-fold changes were considered in further analyses (bolded).

For a species-level determination of the selected OTUs, we performed a nucleotide BLAST search. The OTU AJ243965.1.1512 was identified as *Streptococcus mutans* with 100% query coverage and 100% percentage identity. The lineage report, providing an overview of the taxonomic levels of the hits, found 92 hits out of 100 as *S. mutans*, confirming the species. For OTU FJ470585.1.1487, *Parvimonas micra* was identified as the species with 100% query coverage and 99.77% identity. However, in the lineage report, only four out of 104 hits were for *P. micra*, and most hits failed to categorize it under any genus, thus listing it as an uncultured bacterium or organism. Similarly, OTU JQ451131.1.1383 was identified as *Leptotrichia* genus with 100% query coverage and 100% identity, but no information was available for the species level. Lineage report also identified only six out of 105 hits as uncultured *Leptotrichia* species, while the rest remained as an unidentified bacterium.

### Independent associations between selected OTUs and caries or fillings

We further scrutinized the associations between the selected OTUs and CF. To identify potential determinants of CF, we evaluated age, sex, household income, tooth brushing frequency, Silness-Löe plaque index, Baltic Sea Diet Score, and sucrose intake (E%) in conjunction with the OTUs. These data were available for 389 children. Through the backward stepwise regression, we found that OTUs of *Streptococcus and Parvimonas*, age, and plaque index were independent determinants of CF (**Supplementary Table 2**).

The associations of *Streptococcus* AJ243965.1.1512 and *Parvimonas* FJ470585.1.1487, as dichotomous and categorical variables, with the presence of CF, were tested with logistic regression in crude and adjusted models **(Table 3)**.

**Table 3:** Associations of selected OTUs with CF.

|  |  | With CF |  | Without CF |  | OR<sub>crude</sub> | 95% CI |  | OR<sub>adjusted</sub> | 95% CI |  |
| --- | --- | --- | --- | --- | --- | --- | --- | --- | --- | --- | --- |
|  |  | n | % | N | % |  |  |  |  |  |  |
| *Dichotomous* | ***Streptococcus* AJ243965.1.1512** |  |  |  |  |  |  |  |  |  |  |
|  | Absence | 128 | (52.2) | 117 | (47.8) | Reference |  |  |  |  |  |
|  | Presence | 125 | (86.8) | 19 | (13.2) | **6.01** | **3.49** | **10.36** | **5.81** | **3.291** | **10.26** |
|  | ***Parvimonas* FJ470585.1.1487** |  |  |  |  |  |  |  |  |  |  |
|  | Absence | 147 | (72.8) | 55 | (27.2) | Reference |  |  |  |  |  |
|  | Presence | 106 | (56.7) | 81 | (43.3) | **0.49** | **0.32** | **0.75** | **0.55** | **0.35** | **0.88** |
| *Categorical* |  | With CF |  | Without CF |  | OR | 95% CI |  | OR<sub>adjusted</sub> | 95% CI |  |
|  | ***Streptococcus* AJ243965.1.1512** | n | % | N | % |  |  |  |  |  |  |
|  | Absence | 128 | (52.2) | 117 | (47.8) | Reference |  |  |  |  |  |
|  | Low abundance | 58 | (79.6) | 15 | (20.4) | **3.53** | **1.90** | **6.57** | **3.31** | **1.73** | **6.33** |
|  | High abundance | 67 | (94.4) | 4 | (5.6) | **15.31** | **5.41** | **43.30** | **15.50** | **5.36** | **44.83** |
|  | ***Parvimonas* FJ470585.1.1487** |  |  |  |  |  |  |  |  |  |  |
|  | Absence | 147 | (72.8) | 55 | (27.2) | Reference |  |  |  |  |  |
|  | Low abundance | 64 | (67.4) | 31 | (32.6) | **0.77** | **0.46** | **1.31** | **0.86** | **0.48** | **1.55** |
|  | High abundance | 42 | (45.7) | 50 | (54.3) | **0.31** | **0.19** | **0.53** | **0.36** | **0.20** | **0.64** |

Frequencies of OTUs and their associations with CF in crude and adjusted models for 389 children. Associations of selected OTUs of *Streptococcus* and *Parvimonas* with CF are presented as numbers (frequencies) and odds ratios (ORs) with 95% confidence intervals (CIs) using dichotomous (absence, presence) and categorical (absence, low abundance, high abundance) variables for these OTUs.
Crude model: no adjustments, adjusted model: *Streptococcus* /*Parvimonas* OTU, age, and Silness-Löe plaque index

When using OTUs as dichotomized variables, children with *Streptococcus* AJ243965.1.1512 had a 6 times higher risk of having CF and children with *Parvimonas* FJ470585.1.1487 had a 51% lower risk of having CF compared to children without this OTU. Adjusting for the other OTU, age, and plaque index had only a weak effect on these associations, as children with a *Streptococcus* AJ243965.1.1512 showed a 5.8 times higher risk of having CF and children with a *Parvimonas* FJ470585.1.1487 had a 45% lower risk of having CF compared to children with absence of this OTU. We further assessed the individual and combined associations of these OTUs with having CF and found that when both the *Parvimonas* FJ470585.1.1487 and *Streptococcus* AJ243965.1.1512 were present, the risk of having CF was still elevated but not as high as when the OTU of *Streptococcus* was present alone, without the novel OTU of *Parvimonas* (**Supplementary Table 3).**

In the categorical analysis, compared to children with absence of *Streptococcus* AJ243965.1.1512, the low and high abundance groups showed a 3.5- and a 15.3-times higher risk of having CF in the crude model, respectively (**Table 3**). For *Parvimonas* FJ470585.1.1487, when children with absence of this OTU were compared to those with high abundance, they showed a 69% lower risk of having CF in the crude model. Adjusting for other covariates had only minor impact on these associations.

## Discussion

This study in a general population of children employed 16S rRNA amplicon sequencing to characterize differences in saliva microbiota between children with and without CF. In total, 80 OTUs differed between children with and without CF, with OTUs with the largest differences between the groups further scrutinized. A novel OTU of *Parvimonas* was inversely and independently associated with the risk of having CF, whereas a positive, independent association was observed between an OTU of *Streptococcus* and CF risk. If these microbes co-occurred, *Parvimonas* seemed to mitigate the impact of *Streptococcus* on CF risk.

The results of our nucleotide BLAST search indicated that the novel OTU of *Parvimonas* that inversely associated with CF may be a strain of the *Parvimonas micra* species, which has been mainly considered a periodontal pathogen [32]. More details of the *Parvimonas* taxa and its association with caries are required to infer its role in caries development and prediction.

We observed that a certain *Streptococcus* taxon, most likely a strain of *S. mutans,* demonstrated the strongest positive association with the risk of having CF. *Streptococcus mutans* is a major contributor to the development of caries [33]. However, some *Streptococcus* species, such as *S. salivarius* and *S. mitis,* have been found to protect against caries through pH alkali-generating pathways [34].

We observed compositional differences in a certain *Leptotrichia* taxon that showed a lower abundance in children with CF, but the association was not independent. *Leptotrichia* species are gram-negative, anaerobic rods commonly found in dental plaque [35] and possess high saccharolytic potential [18]. Previously, *Leptotrichia* has been associated positively with the risk of having caries in children under five years [18, 36]. Taken together, our and others results imply that the association of *Leptotrichia* with the risk of having CF depends on the presence of other taxa, and likely on the age, sex, and socioeconomic status of the children. Therefore, further studies are warranted to provide evidence for the putative role of *Leptotrichia* in the development and prediction of caries.

The findings of our earlier study suggested that saliva microbiota composition is sex-specific in adolescents [30], but in the present study among children aged 7-9 years, we observed only one compositional difference between girls and boys. Interestingly, we found that the *TM7x* genus was among the top genera regardless of having CF in boys but not in girls. *TM7x* is a group of ultra-small bacteria in the newly classified “Candidate Phyla Radiation” group [37, 38], and is associated with periodontal diseases such as gingivitis and periodontitis in adults [37, 39]. In addition, the *TM7x* genus has been associated with chronic inflammatory diseases such as juvenile idiopathic arthritis in children and insulin resistance in adults [40, 41]. Hormonal changes during puberty are likely to modify oral microbiota composition [42, 43], which may partly explain why only minor differences in the abundance of saliva bacteria were found between sexes in our population of prepubertal children.

A major strength of the present study is the opportunity to examine a large general population of children with comprehensive data on oral health. Even though dental caries is caused by plaque residing on tooth surfaces, we collected stimulated saliva samples, which is more representative of the oral cavity. Saliva sample collection is also non-invasive; thus, it is easier and more practical to collect than plaque in children. Stimulated saliva has also been shown to result in a higher bacterial count and diversity compared to the collection of unstimulated saliva samples in some but not all studies [44, 45]. We found the associations of *Parvimonas* and *Streptococcus* OTUs with the risk of having CF in a cross-sectional setting, which relies on a single measurement, thus limiting extrapolation of these findings. We did not use international caries diagnostics methods such as International Caries Detection and Assessment System (ICDAS) [46], which can also be considered a limitation of our study. However, our oral health examinations were performed between 2007 and 2009 according to a predefined protocol commonly used for annual oral health examinations in Finland. Additionally, the microbiota analyses did not produce data on the species level, which would have provided more in-depth information on potential microbes associated with having CF, particularly *Parvimonas*. However, the nucleotide BLAST search of the OTUs provided information up to the species level for *Streptococcus* and *Parvimonas*. In this study, we focused on most abundant taxa, and the possibility for other rare OTUs which can have strong associations with CF cannot be ruled out. Lastly, the relative nature of our microbiota data is a limitation. For validating our findings, quantifying the OTUs would be needed. In future studies, shotgun metagenomic sequencing and quantitative analyses (qPCR) should be employed to overcome these limitations and achieve strain-level information [47].

## Conclusions

The present study provides evidence of a novel *Parvimonas* OTU FJ470585.1.1487 that is inversely associated with the risk of having CF. We also showed that a specific OTU of *Streptococcus*, which is likely *S. mutans,* is positively associated with CF. Interestingly, the *TM7x* genus was among the top genera in boys but not in girls, regardless of having CF. Further studies, particularly ones with a longitudinal design and advanced metagenomic sequencing techniques, are warranted to determine the role and mechanism of the novel *Parvimonas* species in the development and prediction of caries.

## List of Abbreviations

CF: Caries or fillings
PANIC: Physical Activity and Nutrition in Children
OTU: operational taxonomic unit
BMI: body mass index
BMI-SDS: body mass index standard deviation score
OR: odds ratio
CI: confidence interval
BSDS: Baltic Sea Diet Score

## Declarations

### Ethical approval and consent to participate

The Research Ethics Committee of the Hospital District of Northern Savo approved the study protocol in 2006 (Statement 69/2006). The parents or caregivers of the children gave their written informed consent, and the children provided their assent to participation.

### Consent for publication

Not applicable

## Availability of data and materials

The datasets generated and/or analyzed during the current study are available in the European Genome-phenome Archive (EGA) repository (https://ega-archive.org/datasets/EGAD50000000989), and full access to the repository could be arranged on request from the corresponding author. The data are not publicly available due to restricted consent from the participants.

## Author’s contributions

VFK, NA, TAL, HV conceptualized the study. VFK, NA, AME, ALS, MN, and TAL curated the data for the study. NA and HV designed the microbiota methodologies and performed the analysis and data validation used in the study. NA and VFK wrote the original draft. All authors critically revised the manuscript. HV and TAL supervised the study and were responsible for funding acquisition.

## Acknowledgements

We are grateful to all children and their parents participating in the PANIC Study. We are also indebted to the research group and personnel of the PANIC study for their invaluable contribution throughout the study.

