## Supplementary material for "A novel *Parvimonas* OTU is inversely associated with having caries or fillings in children"

**Supplementary Table 1:** Composition percentages at Phylum level between children with caries and without caries in all, girls and boys separately

| **Taxa** | **All together - Abundance percentage (n=400)** | | **Girls - Abundance percentage (n=180)** | | **Boys - Abundance percentage (n=220)** | |
| --- | --- | --- | --- | --- | --- | --- |
|  | **With CF**  **(n=261)** | **Without CF**  **(n=139)** | **With CF (n=119)** | **Without CF (n=61)** | **With CF**  **(n=142)** | **Without CF**  **(n=78)** |
| **Bacillota** | 68.86 | 68.66 | 68.94 | 68.29 | 68.79 | 68.94 |
| **Actinomycetota** | 15.87 | 15.22 | 15.53 | 15.24 | 16.15 | 15.20 |
| **Bacteroidota** | 8.97 | 8.52 | 9.16 | 8.92 | 8.81 | 8.21 |
| **Pseudomonadota** | 2.65 | 3.62 | 2.56 | 3.93 | 2.72 | 3.38 |
| **Patescibacteria** | 2.08 | 2.24 | 2.11 | 1.93 | 2.05 | 2.48 |
| **Fusobacteriota** | 1.49 | 1.66 | 1.62 | 1.61 | 1.39 | 1.70 |
| **Other** | 0.05 | 0.05 | 0.05 | 0.04 | 0.05 | 0.05 |
| **Total** | **99.97** | **99.97** | **99.97** | **99.96** | **99.96** | **99.96** |

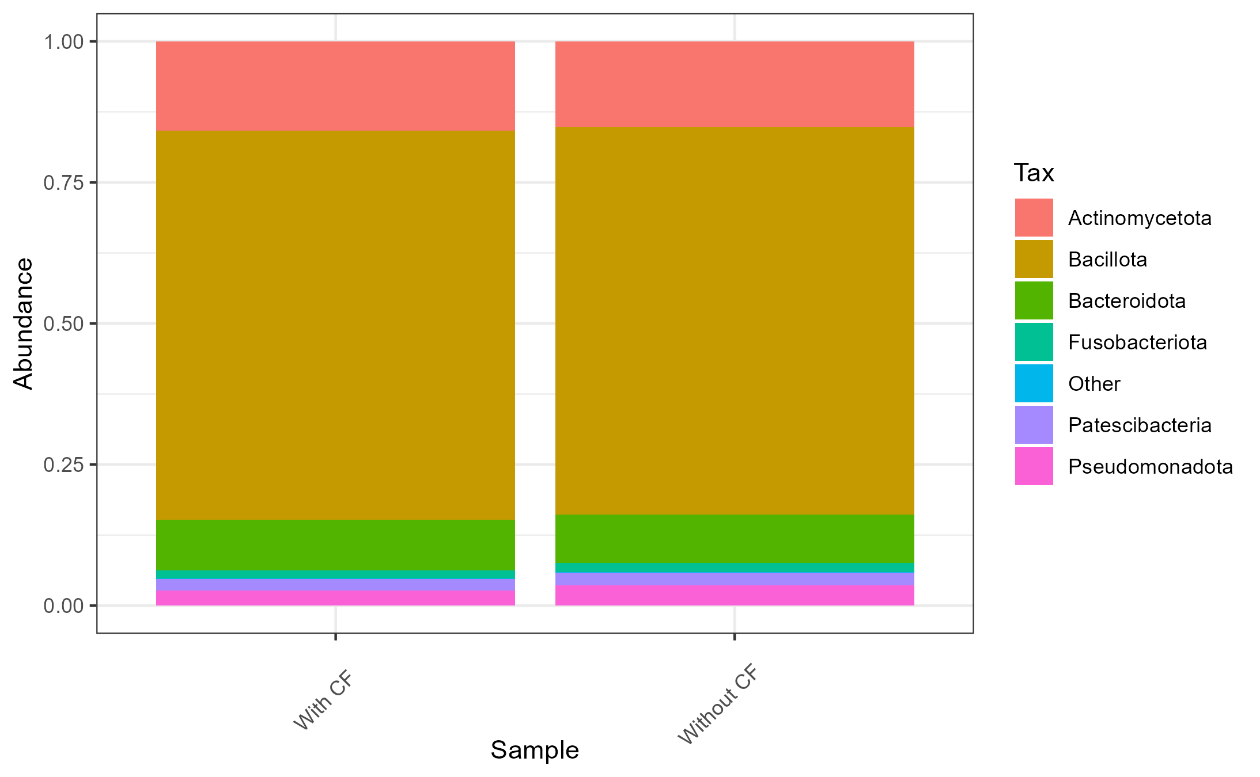

**Supplementary Figure 1:** Composition at Phylum level between children with caries and without caries in all, boys and girls separately: Panel A) Altogether, B) Girls and C) Boys.

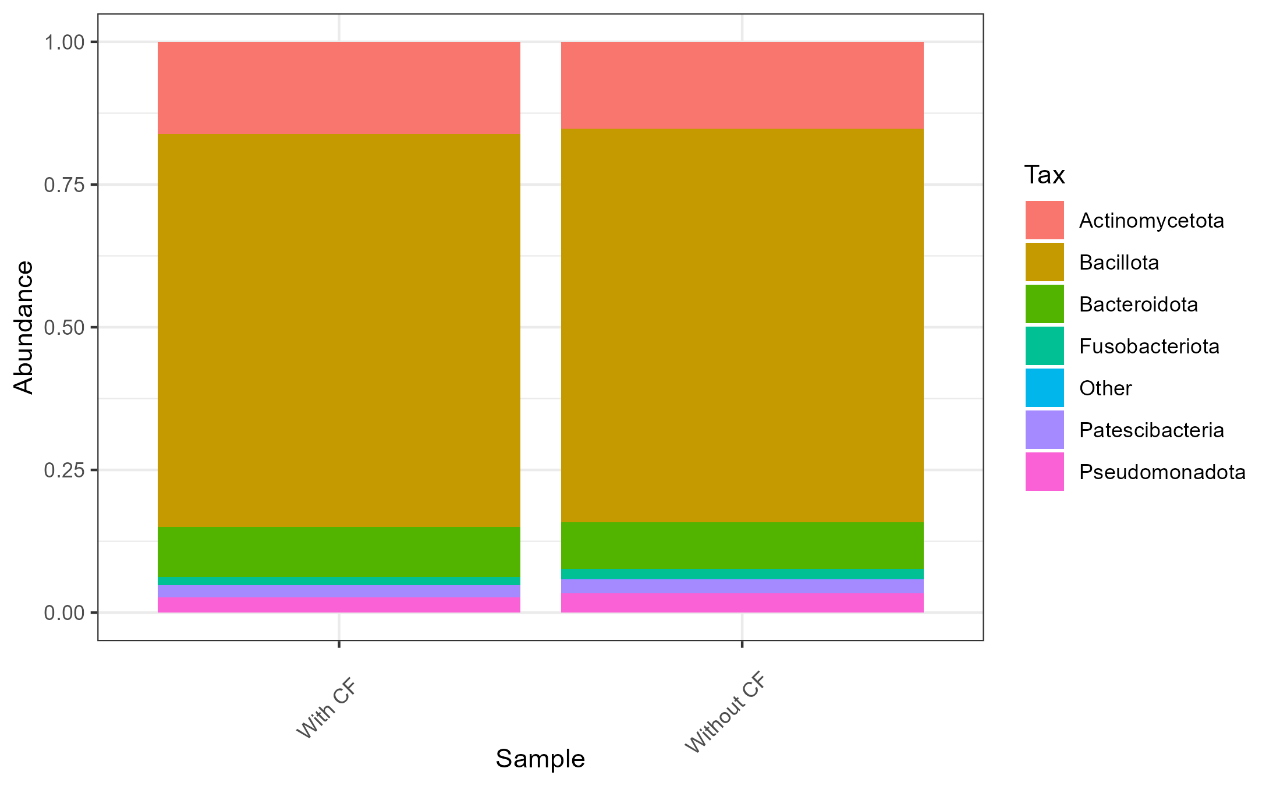

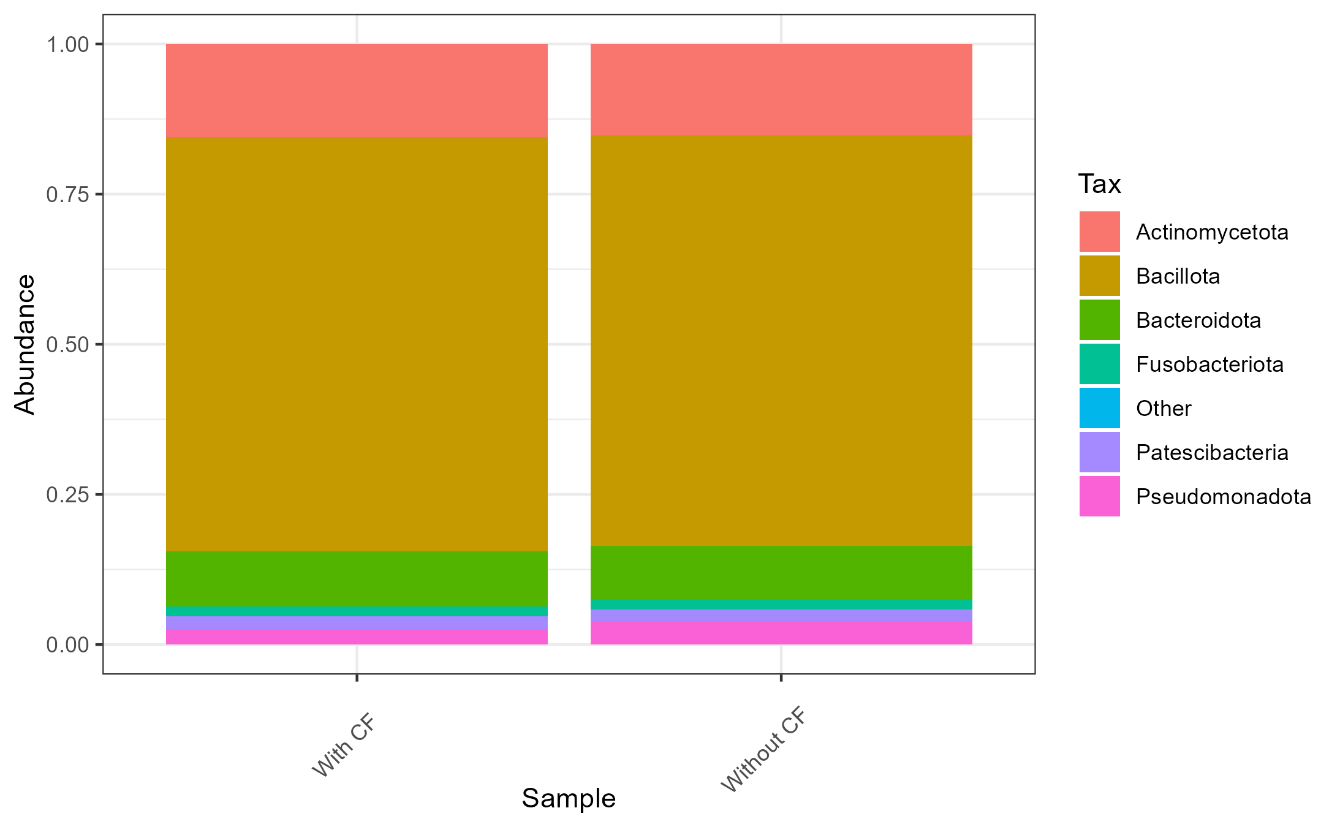

Panel C – Boys

Panel A – Altogether

Panel B – Girls

| **Supplementary Table 2. Variables remaining in the model after backward stepwise logistic regression with Wald method. For each variable, B coefficient, SE, OR and 95% CI for OR is reported.** | | | | | | | | |
| --- | --- | --- | --- | --- | --- | --- | --- | --- |
| **Model** | **B** | **SE** | **Wald** | **df** | **p-value** | **OR** | **95% CI for OR** | |
|  |  |  |  |  |  |  | **Lower** | **Upper** |
| Streptococcus (0/1) | 1.503 | 0.306 | 24.153 | 1 | <.001 | 4.49 | 2.47 | 8.18 |
| Parvimonas (0/1) | -0.609 | 0.257 | 5.594 | 1 | 0.018 | 0.54 | 0.33 | 0.90 |
| The Silness-Löe plaque index | 0.158 | 0.048 | 10.712 | 1 | 0.001 | 1.172 | 1.07 | 1.29 |
| Age, years | -0.682 | 0.343 | 3.958 | 1 | 0.047 | 0.51 | 0.26 | 0.99 |
| Constant | 4.996 | 2.635 | 3.595 | 1 | 0.058 | 147.76 |  |  |
| Variables included: Leptotrichia (0/1), Streptococcus (0/1), Parvimonas (0/1), sex, household income, Baltic Sea Diet Score, Silness-Löe plaque index, age, sucrose intake E%, tooth brushing frequency | | | | | | | | |

| **Supplementary Table 3. Independent and combined associations of Streptococcus and Parvimonas OTUs with having caries or fillings (CF)** | | | | | | | | | | |
| --- | --- | --- | --- | --- | --- | --- | --- | --- | --- | --- |
|  | **With CF**  **(n=253)** | | **Without CF**  **(n=136)** | | **OR_crude_** | **95% CI** | | **OR_adjusted_** | **95% CI** | |
|  | **n** | **%** | **n** | **%** |  |  |  |  |  |  |
| Neither OTU | 62 | (24.5) | 48 | (35.3) | REF =1 |  |  | REF=1 |  |  |
| Only Streptococcus OTU | 85 | (33.6) | 7 | (5.1) | 9.40 | 3.99 | 22.17 | 9.42 | 3.90 | 22.72 |
| Only Parvimonas OTU | 66 | (26.1) | 69 | (50.7) | 0.74 | 0.45 | 1.23 | 0.67 | 0.40 | 1.15 |
| Both OTUs | 40 | (15.8) | 12 | (8.8) | 2.58 | 1.22 | 5.45 | 2.56 | 1.19 | 5.53 |
| Crude model: no adjustments, adjusted model: age and Silness-Löe plaque index | | | | |  |  |  |  |  |  |
